# Structural basis of nucleotide selectivity in pyruvate kinase

**DOI:** 10.1101/2024.06.25.600556

**Authors:** Atsushi Taguchi, Ryosuke Nakashima, Kunihiko Nishino

## Abstract

Nucleoside triphosphates are indispensable in numerous biological processes, with enzymes involved in their biogenesis playing pivotal roles in cell proliferation. Pyruvate kinase (PYK), commonly regarded as the terminal glycolytic enzyme that generates ATP in tandem with pyruvate, is also capable of synthesizing a wide range of nucleoside triphosphates from their diphosphate precursors. Despite their substrate promiscuity, some PYKs show preference towards specific nucleotides, suggesting an underlying mechanism for differentiating nucleotide bases. However, the thorough characterization of this mechanism has been hindered by the paucity of nucleotide-bound PYK structures. Here, we present crystal structures of *Streptococcus pneumoniae* PYK in complex with four different nucleotides. These structures facilitate direct comparison of the protein-nucleotide interactions and offer structural insights into its pronounced selectivity for GTP synthesis. Notably, this selectivity is dependent on a sequence motif in the nucleotide recognition site that is widely present among prokaryotic PYKs, particularly in Firmicutes species. We show that pneumococcal cell growth is significantly impaired when expressing a PYK variant with compromised GTP and UTP synthesis activity, underscoring the importance of PYK in maintaining nucleotide homeostasis. Our findings collectively advance our understanding of PYK biochemistry and prokaryotic metabolism.

## INTRODUCTION

Glycolysis is an omnipresent metabolic pathway classically described as an anaerobic process that produces a net gain of two ATP molecules from one glucose molecule. The last step of glycolysis is catalyzed by pyruvate kinase (PYK), which produces ATP and pyruvate via a phosphoryl transfer reaction from phosphoenolpyruvate (PEP) to ADP (1). This forward reaction is essentially irreversible in cells and makes PYK a key regulator of metabolic flux through this pathway. Recently, PYK has garnered renewed interest due to its presumed role in altering glucose metabolism in cancer cells, known as the Warburg effect (2, 3). PYK also plays a central role in bacterial metabolism, making it a promising target for novel antibiotic development against multidrug-resistant bacteria (4).

Over the years, PYKs from various species have been structurally characterized to gain molecular insight into their regulatory mechanism that could aid in developing therapeutic interventions. The majority of characterized PYKs are homotetramers allosterically regulated by sugar phosphate intermediates in the metabolic pathway and/or amino acids, which allows flux control in response to changes in the cellular environment (5–7). Unlike some allosteric enzymes that undergo significant conformational changes at the active site upon effector binding, PYK exhibits only subtle alternations at the active site. Instead, PYK displays a rocking motion mechanism in which effector binding triggers a rigid body rotation of each protomer and the rearrangement of inter-protomer interactions (8–10).

Although the exact mechanism of allosteric regulation varies among PYKs and remains an active area of research, it is generally accepted that this rearrangement of inter-protomer interactions stabilizes PYK in its active state conformation and increases substrate affinity.

While many aspects of PYK activity have extensively been studied, the structural basis of nucleotide differentiation has received little attention despite its importance in defining PYK function. Although ADP is commonly regarded as the primary PYK substrate because of its role in energy generation, it has long been recognized that PYKs can utilize other NDPs as phosphate acceptors and in some cases prefer them over ADP (Figure 1A). For instance, GDP has been reported to be the best phosphate acceptor for *Escherichia coli* PykF and *Streptococcus mutans* PYK (11, 12). In the parasite *Plasmodium falciparum*, the broad NTP synthesis activity of PYK is critical for organelle maintenance under physiological conditions (13). These examples likely represent the significant role of PYK in maintaining nucleotide homeostasis in many organisms and suggest that PYKs possess structural features that allow nucleotide differentiation. However, our understanding of the nucleotide binding site remains limited because of the lack of PYK structures in complex with nucleotides other than ADP or ATP in the active site (1). Furthermore, the existing ADP/ATP-bound PYK structures contain few interactions between the adenine base and the active site residues, prompting questions about the mechanism underlying nucleotide selectivity in PYKs (10, 14).

**Figure 1:**
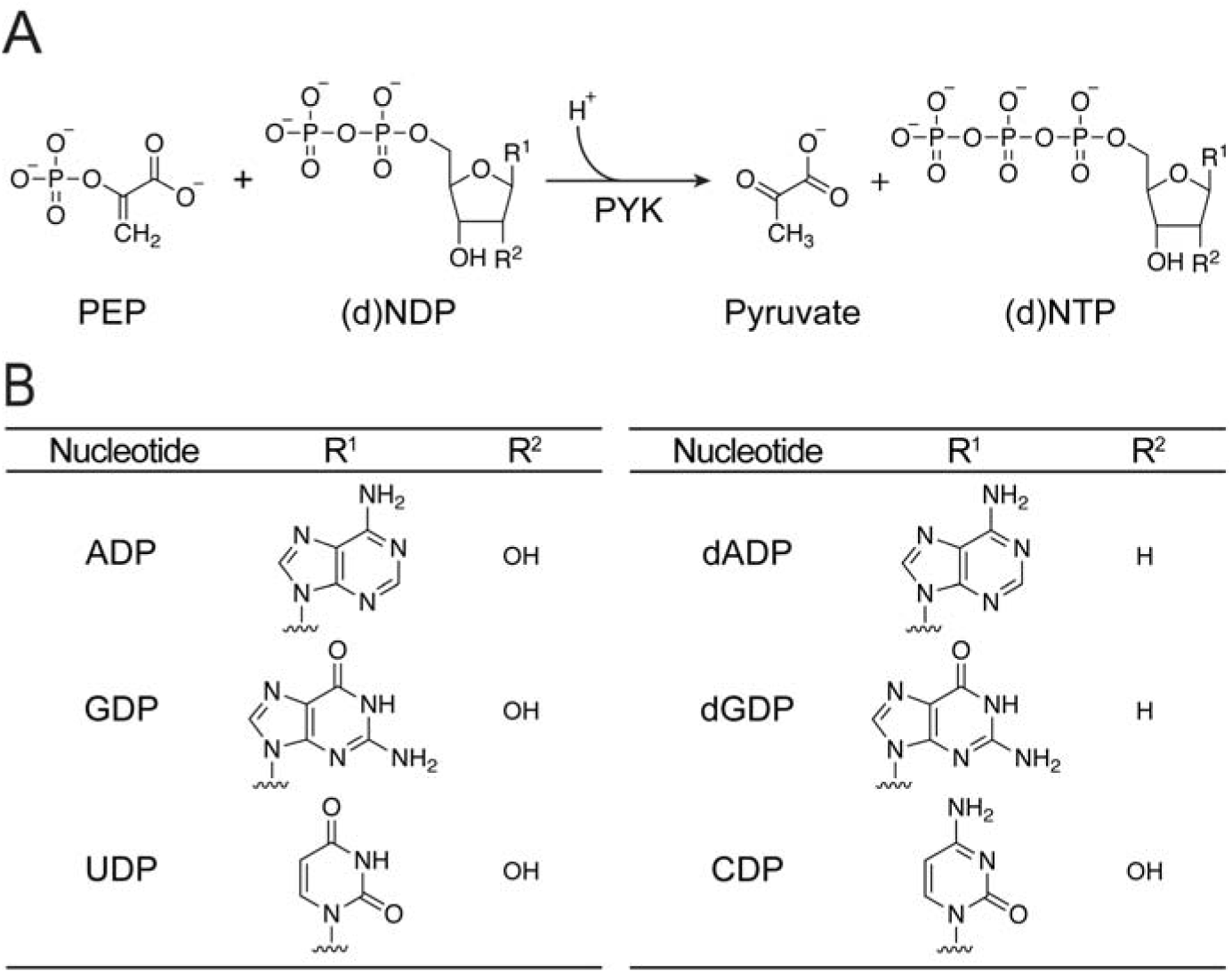
PYK synthesizes nucleoside triphosphates from their diphosphate precursors. (A) Schematic of the PYK enzymatic reaction that yields pyruvate and (d)NTP. *Sp*PYK requires potassium and magnesium ions to catalyze this reaction (18). (B) Chemical structures of the nucleotide substrates that were used to assess *Sp*PYK activity in this study.

In this study, we sought to address this question through structural determination of nucleotide-bound PYK from *Streptococcus pneumoniae*. *S. pneumoniae* is an opportunistic human pathogen that can cause diseases ranging from mild respiratory tract infections to severe conditions such as meningitis and pneumonia, and it is also one of the leading pathogens for mortalities associated with antimicrobial resistance (15, 16). Therefore, elucidating the functional and structural properties of essential enzymes like PYK, which represent potential targets for new antimicrobials, is of utmost interest (17). Here, we present the first high-resolution crystal structures of PYK in complex with GDP or UDP, enabling direct comparison of the nucleotide recognition site with the ADP-bound structure.

Our findings reveal GDP as the preferred substrate for *S. pneumoniae* PYK (*Sp*PYK), and we provide a structural rationale for this substrate preference by identifying a conserved sequence motif in the active site shared among a subset of prokaryotic PYKs. By introducing alanine mutations in this motif, we succeeded in generating a *Sp*PYK variant with altered nucleotide selectivity, allowing us to probe the impact of NTP synthesis activity by *Sp*PYK on cell growth. This work fills a gap in our understanding of the functional properties of PYK and sheds light on its role in nucleotide metabolism.

## RESULTS

### GDP is the preferred substrate for *Sp*PYK

Previously, we demonstrated that *Sp*PYK is allosterically activated by the glycolytic intermediate fructose 1,6-bisphosphate (FBP); however, the scope of its nucleotide substrate remained unexplored (18). To investigate its nucleotide preference, we measured the steady-state kinetics of *Sp*PYK in the presence of FBP with select NDPs and dNDPs (Figure 1B). We detected PYK activity for every nucleotide tested, but we observed a considerable difference in the level of PYK activity depending on the nucleotide substrate (Table 1). GDP was the best phosphate acceptor with a K_m_ value of 0.027 ± 0.002 mM, which was ∼50-fold lower than that of ADP. UDP was a slightly better *Sp*PYK substrate compared with ADP based on the catalytic efficiency inferred from the k_cat_/K_m_ values. CDP was the least favored phosphate acceptor among the NDPs, with a k_cat_/K_m_ value approximately 300-fold lower than that of GDP. These results are consistent with *Sp*PYK having a mechanism for distinguishing nucleotide base structure and preferentially using GDP to catalyze PEP to pyruvate conversion. A similar trend in nucleotide utilization was reported for *S. mutans* PYK, which is activated by glucose 6-phosphate instead of FBP, indicating that streptococcal PYKs may broadly share this nucleotide preference even when they are activated by different sugar-phosphates (12). Furthermore, we observed a 4-6 fold decrease in k_cat_/K_m_ values when dNDPs were used instead of their NDP counterparts, indicating that the ribose 2’ hydroxyl group is also involved in protein-ligand interactions but is not strictly required for substrate recognition by *Sp*PYK.

**Table 1:** Kinetic properties of *Sp*PYK with respect to nucleoside diphosphate substrates. All data are mean ± SEM from experiments conducted in triplicate. *Sp*PYK displays a hyperbolic kinetic profile in the presence of its activator FBP, and kinetic parameters were obtained by fitting the data into the Michaelis-Menten equation.

| Nucleotide | $K_m$ (mM) | $k_{cat}$ ($s^{-1}$) | $k_{cat}/K_m$ |
| --- | --- | --- | --- |
| ADP | $1.32 \pm 0.07$ | $353.1 \pm 7.4$ | 268 |
| GDP | $0.027 \pm 0.002$ | $280.0 \pm 4.0$ | 9334 |
| UDP | $0.76 \pm 0.03$ | $311.2 \pm 4.3$ | 410 |
| CDP | $1.95 \pm 0.19$ | $63.7 \pm 1.4$ | 33 |
| dADP | $1.91 \pm 0.08$ | $119.1 \pm 1.9$ | 62 |
| dGDP | $0.16 \pm 0.01$ | $222.4 \pm 2.9$ | 1588 |

### The sugar-phosphate binding site in PYK is highly conserved

To gain a better understanding of how nucleotides are recognized by *Sp*PYK, we carried out vapor diffusion crystallization trials with various nucleotides in the presence of FBP and the enolpyruvate analog oxalate to stabilize the protein in its active conformation. We successfully obtained crystals that diffracted to ∼2 Å under conditions that included ATP, ADP, GDP, or UDP, and subsequently solved the *Sp*PYK structures with each of these nucleotides (Figure 1A, Table S1). While the nucleotide binding sites in the ADP- and GDP-bound structures were fully saturated, ATP was found to be present in only half of the protomers in its structure (Figure S1). For UDP, we solved two crystal structures with two, three or four nucleotides per tetramer (Figure S1). The absence of nucleotides in some PYK protomers, which has also been reported in previous structural studies for other PYKs, may be due to forces incurred during crystal lattice formation (9, 14).

In the protomers containing the desired nucleotides, the bound nucleotide and oxalate were present in the active site located in a cleft formed between the A domain with a (β/17)_8_ barrel fold and the mobile B domain (Figure 2A). FBP is bound to the C domain effector binding site proximal to the protomer interface and locks *Sp*PYK in its active state. We previously identified active site residues and cations (K^+^ and Mg^2+^) that mediate PEP binding in the PEP-bound *Sp*PYK structure (18). These residues and cations were also found to interact with the oxalate and the ATP γ-phosphate in the ATP-bound structure, which mimics the enzyme state after the phosphoryl transfer reaction (Figure 2B). The ATP-bound structure contains a second Mg^2+^ that is stabilized through an octahedral coordination involving nucleotide phosphates as well as three water molecules that form hydrogen bonds with Thr306 and Ser340 in the A domain and Asp153 in the B domain, and this Mg^2+^ is also retained in the NDP structures that lack the γ-phosphate (Figure 2C). Contacts observed in the active site involving the 17- and β-phosphates are well-preserved across our solved structures, which include a water molecule bridge between the Gly185 amide and the nucleotide β-phosphate. The ribose 2’ and 3’ hydroxyl groups are stabilized through interactions with Lys184, which explains the difference in substrate affinity observed between NDPs and dNDPs. Overall, the interactions between *Sp*PYK and the ribose-phosphate moiety resemble those reported in other PYKs, highlighting the highly conserved nature of the catalytic site (9, 14, 19).

**Figure 2:**
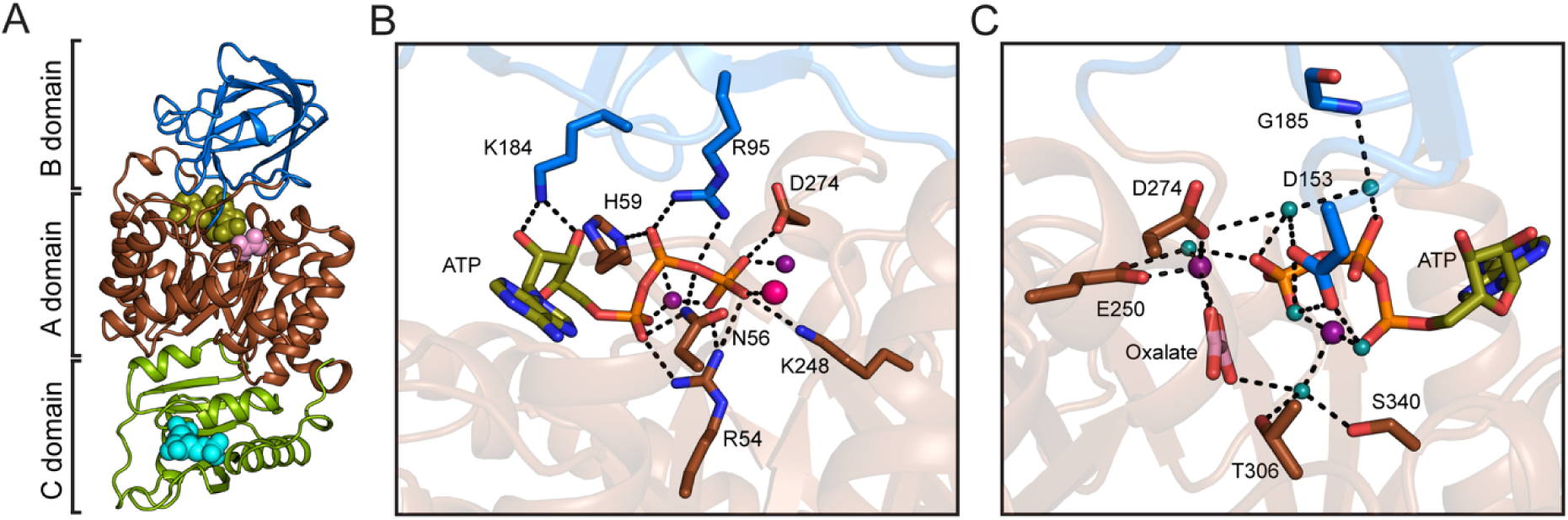
Coordination of the ATP sugar-phosphate moiety by *Sp*PYK. (A) Crystal structure of the ATP-bound *Sp*PYK protomer (PDB: 8XW6). The brown, blue, and light green regions correspond to the A, B, and C domains, respectively. Oxalate (light pink) and ATP (olive) are bound to the active site sandwiched by AB domains. FBP (cyan) is located at the C domain effector binding site. (B) A close-up view of active site residues and cations that directly interact with the sugar-phosphate moiety of ATP. Pink and purple spheres correspond to K^+^ and Mg^2+^ ions, respectively. Oxalate is omitted for clarity. (C) Water molecules are present in the hydrogen bond network that coordinates the ATP sugar-phosphate moiety. Active site residues and cations that interact with water (teal) are shown.

### AIZ2 helix residues are responsible for nucleotide selectivity in *Sp*PYK

Compared with the active site residues responsible for ribose-phosphate recognition, those surrounding the nucleotide base are more variable (18). Our NDP-bound structures provide key insights into how these residues allow *Sp*PYK to distinguish incoming nucleotides. The only *Sp*PYK-nucleotide base interaction seen in the ADP-bound structure is a direct contact between Glu64 at the N6-adenine (Figure 3A). This was in stark contrast to the higher number of guanine-*Sp*PYK contacts identified in the GDP-bound structure (Figure 3B). Glu64 interacts with the guanine at the N1 and N2 positions, with the latter interaction mediated by a water molecule. In addition, Gln65 and Arg68 coordinate the O6-guanine and a water molecule which in turn stabilizes a second water molecule that bridges the N7-guanine with the Leu11 carbonyl and the Asn56 amide. Despite the switch from a purine to a pyrimidine in the UDP-bound structure, these water molecules maintain identical contacts with the *Sp*PYK residues by coordinating the O4-uridine (Figure 3C). UDP-bound protomers in the half-saturated structure also contain a direct interaction between the O4-uridine and Arg68, and these protomers form water-mediated contacts involving Glu64 and Asn345 with N3-uridine and O2-uridine, respectively.

**Figure 3:**
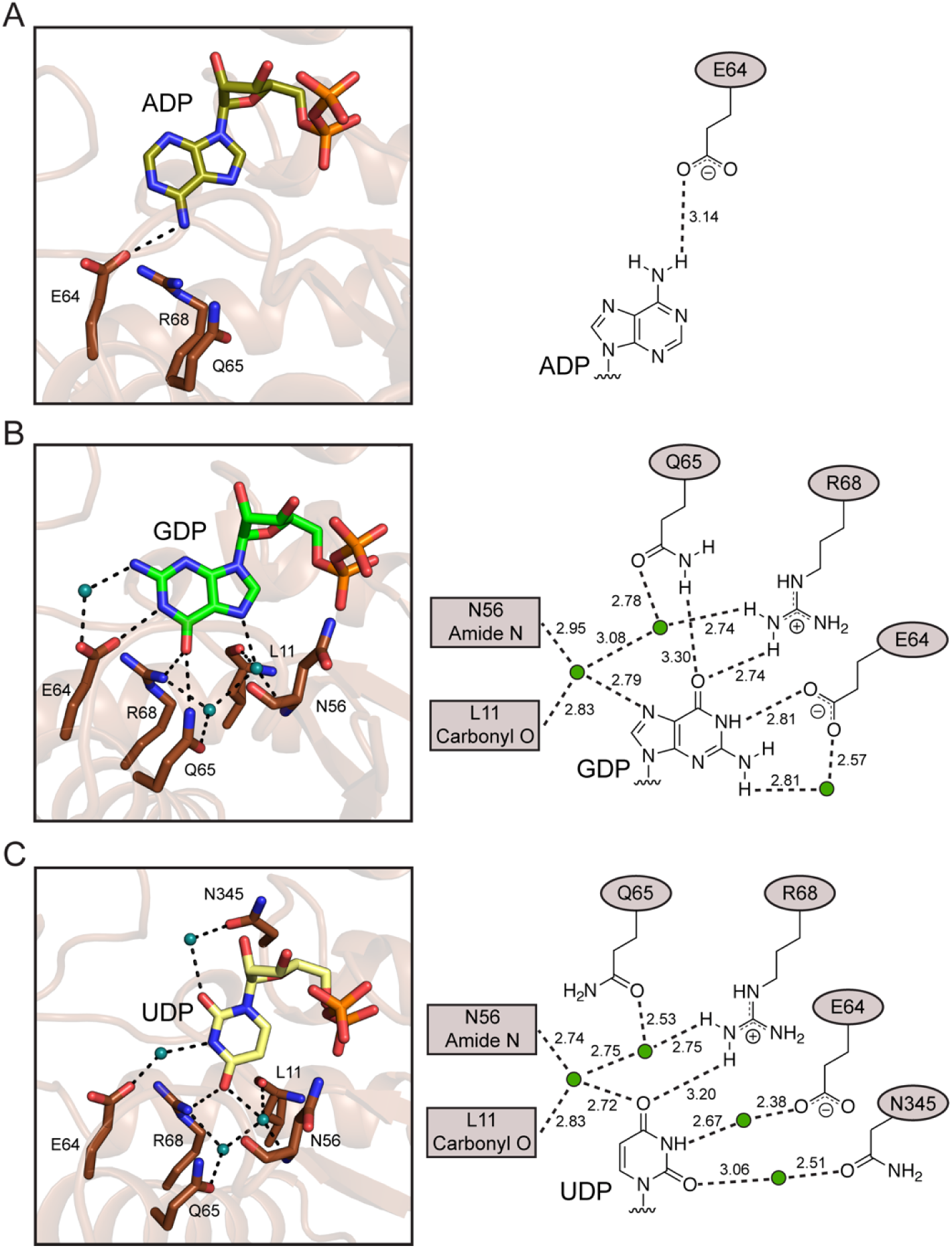
*Sp*PYK can structurally discriminate incoming nucleotide bases. Crystal structures of the ADP-bound (A, PDB: 8XW7), GDP-bound (B, PDB: 8XW8) and UDP-bound (C, PDB: 8XW9) *Sp*PYK active sites are shown in the left panels. Corresponding schematic drawings of putative interactions between the nucleotide base and active site residues along with the observed distances in angstroms are shown in the right panels. Water molecules are depicted as teal spheres.

Some of these interactions were not found in other UDP-bound protomers, implying the flexible nature of *Sp*PYK UDP recognition (Figure S2). Overall, the number of nucleotide base-protein contacts corresponds to the nucleotide preference exhibited by *Sp*PYK, suggesting that it can discriminate among nucleotide substrates through physical interactions.

We were intrigued by the involvement of Glu64, Gln65 and Arg68 in nucleotide base coordination because A172 helix residues have not been implicated in substrate recognition. In other ADP/ATP-bound PYK structures, the adenine ring does not directly interact with the active site residues but rather interacts with water molecules coordinated by residues analogous to Leu11 and Asn56 in our GDP- and UDP-bound *Sp*PYK structures (Figure S3). To better understand the role of nucleotide-interacting residues on the *Sp*PYK A172 helix in nucleotide recognition, we prepared single point mutants with alanine substitutions as well as double (E64AR68A) and triple (E64AQ65AR68A; AAA) mutants, and assessed their catalytic activity using ADP, GDP or UDP as their substrate (Figure 4A, Figure S4). If the interaction between the nucleotide base and the protein were crucial for ADP recognition, we would expect a significant reduction in PYK activity of the E64A mutant with ADP, as Glu64 is the sole residue that directly interacts with adenine. However, we observed only a moderate decrease (∼30%) in activity for this mutant, indicating that ADP coordination at the active site does not require this specific interaction. Unexpectedly, replacing Arg68 with alanine led to a ∼30% increase in PYK activity under our experimental conditions, which implied that this arginine negatively affects ADP binding. While a larger decrease in activity was observed for Q65A and AAA mutants compared to the E64A mutant, these mutants still retained ∼50% of wild-type PYK activity. These findings suggest that *Sp*PYK might adopt a mode of ADP binding similar to what has been reported in other PYKs when specific residues in the A172 helix residues are mutated (Figure S3).

**Figure 4:**
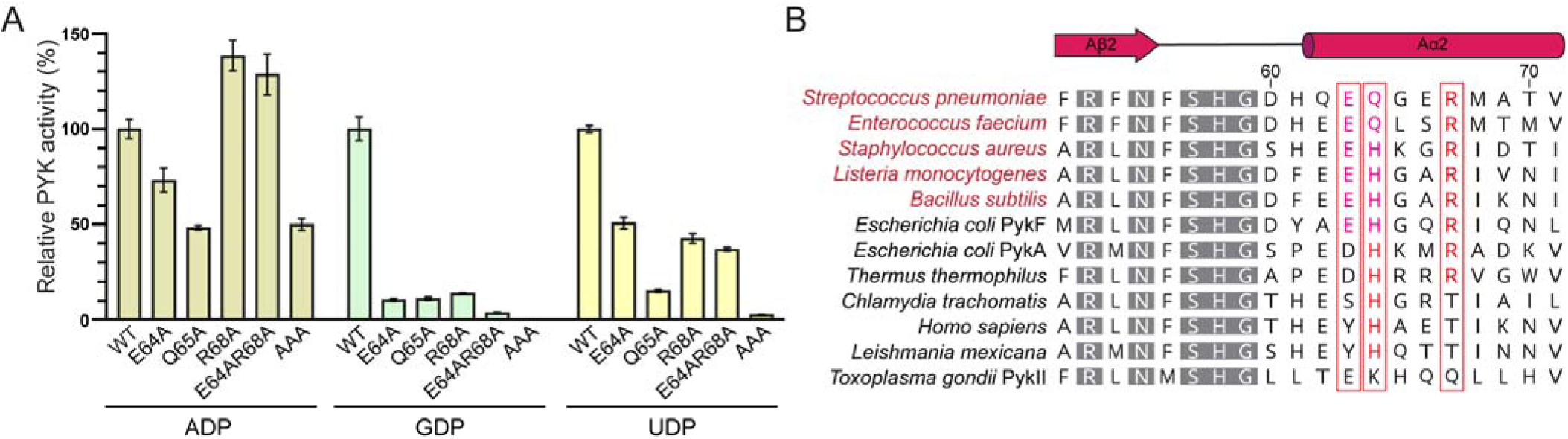
The AIZ2 helix motif confers nucleotide selectivity. (A) *Sp*PYK activity of wild-type (WT) and indicated mutants was measured *in vitro* in the presence of 1 mM ADP, 0.25 mM GDP, or 1 mM UDP to assess the contribution of A172 helix residues towards nucleotide selectivity. Error bars represent mean ± SD from triplicates. The value obtained for the GDP AAA mutant was very small (0.20 ± 0.05%) and is not displayed in the figure. A statistically significant difference (P < 0.001, one way ANOVA with Dunnett’s test, GraphPad Prism 9.0) was observed between the WT sample and mutant samples for all combinations. (B) Sequence alignment of A172 helix residues and upstream residues in representative PYKs. Residue numbers and secondary structures are based on the *Sp*PYK structure. Gray residues are conserved in >80% of the PYK sequences (18). Red boxes denote the location of the E(Q/H)xxR motif, and residues that match this motif are shown in red. This motif is widely present in Firmicutes PYKs, including *E. faecium*, *S. aureus*, *L. monocytogenes* and *B. subtilis*. See Figure S5 for the full sequence alignment.

In contrast, a notable difference in PYK activity was observed when GDP or UDP served as the substrate (Figure 4A). When GDP was used, PYK activity decreased approximately 10-fold upon alanine replacement of any of the three A172 helix residues, and the activity was further reduced in the double mutant. A similar decrease in activity among the mutants was observed for UDP, although the decrease was less pronounced. In both cases, introduction of the triple mutation had the most detrimental effect on PYK activity, with the AAA mutant exhibiting only ∼0.2% and ∼3% of wild-type PYK activity for GDP and UDP, respectively. Taken together, our results indicate that A172 helix residues play a significant role in guanine and uridine recognition, while their contribution towards adenine recognition appears to be relatively minor.

### A**IZ**2 helix motif is conserved among a subset of prokaryotic PYKs

Having demonstrated the importance of A172 helix residues in establishing *Sp*PYK nucleotide selectivity, we next performed a sequence alignment of the A172 helix from a diverse set of prokaryotic and eukaryotic PYKs to evaluate the prevalence of these residues in other species (Figure 4B and Figure S5). Our analysis revealed that the EQxxR motif is primarily found in streptococcal and enterococcal species, whereas the glutamine is replaced by a histidine (EHxxR) in other Firmicutes species. The same sequence motif is also present in one of the two PYK isozymes encoded in certain Proteobacteria species. *E. coli* PykF is one such example, and this isozyme has been reported to prefer GDP as the nucleotide acceptor, which implies that the histidine replacement does not substantially affect GDP binding (11). On the other hand, a glutamate-to-aspartate substitution appears to abolish GDP selectivity as exemplified by *E. coli* PykA and *Thermus thermophilus* PYK (20, 21). Some prokaryotic PYKs (*e.g., Chlamydia trachomatis* PYK) and eukaryotic PYKs lack both glutamate and arginine residues and typically do not demonstrate GDP selectivity (22, 23). A notable exception to this rule is *Toxoplasma gondii* PykII, which demonstrates a clear preference towards GDP despite the absence of the E(Q/H)xxR motif, indicating that this PYK has a different mode of guanine recognition (24). In summary, our sequence analysis suggests that the E(Q/H)xxR motif on the A172 helix is a characteristic feature of prokaryotic PYKs that favor GDP utilization over other NDPs.

### PYK nucleotide preference plays a key role in pneumococcal metabolism

Given the broad conservation of the E(Q/H)xxR motif among prokaryotic PYKs, we wondered whether preferential synthesis of certain NTPs by *Sp*PYK is important in pneumococcal metabolism. We reasoned that characterizing pneumococcal cells expressing the AAA mutant could provide insight into the physiological relevance of *Sp*PYK nucleotide selectivity because this mutant still retains the ability to carry out glycolysis through the use of ADP while its GTP and UTP synthesis activities are greatly reduced. To confirm whether the AAA mutant still catalyzes pyruvate synthesis in cells, we turned to a previously established fosfomycin sensitivity assay (18). Fosfomycin inhibits the cell wall enzyme MurA, which uses PEP to generate cell wall precursors, and cells with impaired PYK activity become resistant to this cell wall antibiotic due to the accumulation of PEP that can outcompete fosfomycin for the MurA active site (18, 25). We introduced the AAA mutation at the native *pyk* locus of an *S. pneumoniae* strain with an IPTG-inducible wild-type *pyk* copy at the ectopic site that facilitates replacement of the essential *pyk* gene. This strain was then plated on an fosfomycin agar plate that allows the growth of a *pyk*(H411A) mutant strain, which is predicted to have lower PYK activity regardless of the NDP substrate because of a mutation in the FBP binding site (Figure 5A). We found that cells expressing the AAA mutant were inviable on the fosfomycin plate, indicating that the glycolytic flow is not severely affected in these cells.

**Figure 5:**
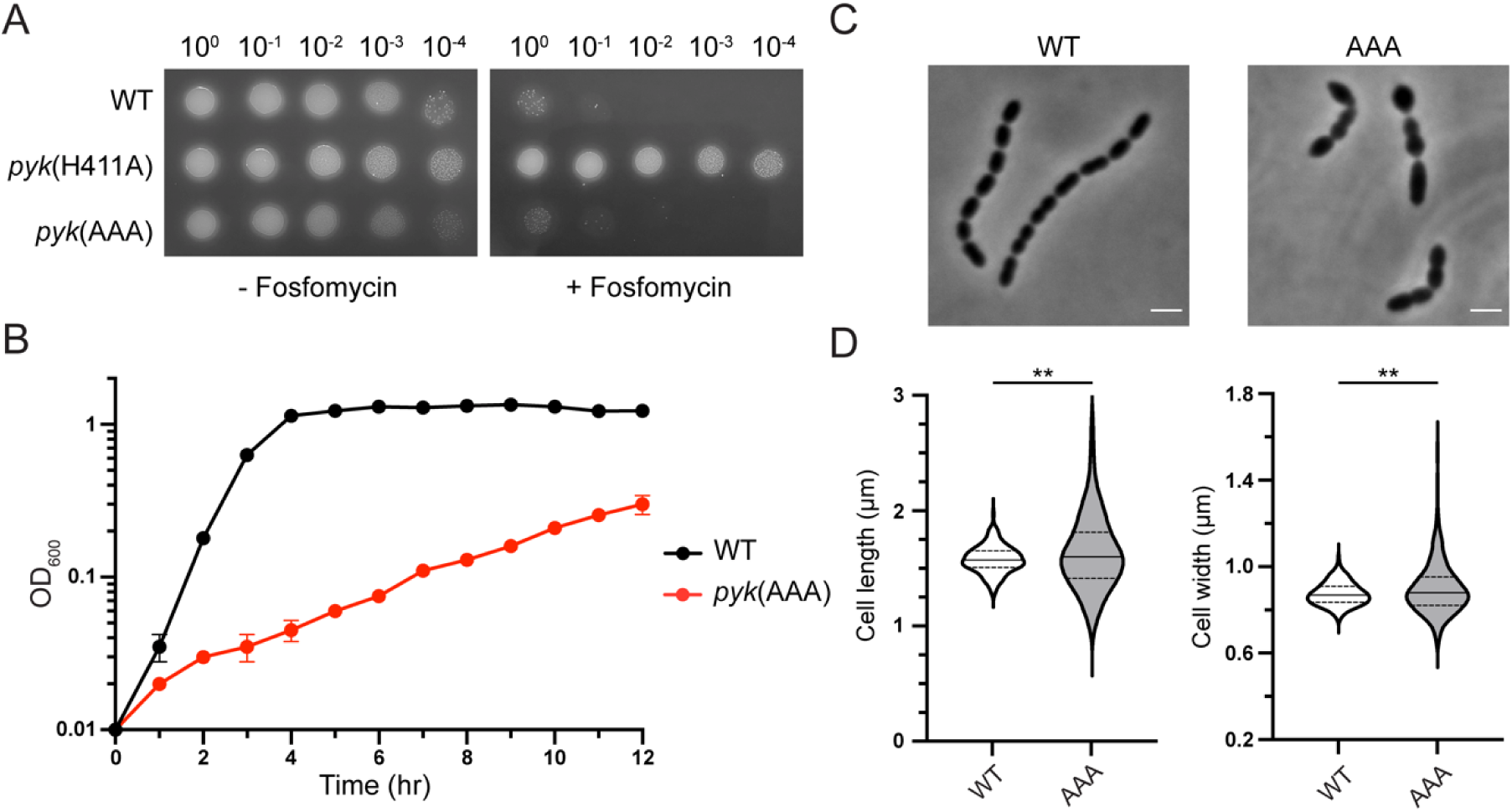
PYK NTP synthesis activity is important for pneumococcal growth. (A) *S. pneumoniae* R6 cells containing the indicated *pyk* gene at the native *pyk* locus were spotted on an agar plate with or without fosfomycin (64 µg/mL) to assess viability. (B) Representative growth curves of *S. pneumoniae* D39 WT and *pyk*(AAA) strains. Error bars represent mean ± SD from duplicates. (C) Phase-contrast images of *S. pneumoniae* D39 WT and *pyk*(AAA) cells. Scale bar: 2 µm. (D) Violin plots of cell length and cell width of D39 WT and *pyk*(AAA) cells. Solid and dotted lines indicate the median and quartile, respectively. **P<0.005 (unpaired t-test). See Supplementary Information for details on the genetic background of R6 and D39 cells used in this study.

We proceeded to assess the effect of expressing the AAA mutant on cell growth and morphology. For these experiments, we deleted the ectopic wild-type *pyk* copy from cells containing the AAA mutation at the native locus to ascertain that any phenotypic differences are attributed to the AAA mutation. A striking difference in growth rate was observed between the wild-type and the AAA mutant; while the wild-type reached stationary phase after 4 hrs post-incubation, the AAA mutant grew extremely slowly and only reached ∼1/4 of the cell density achieved by the wild-type strain 12 hr post-incubation (Figure 5B). Moreover, phase-contrast micrographs of AAA mutant cells revealed heterogeneity in cell size, indicative of defects in cell growth and division that prevent proper cell shape maintenance (Figures 5C and 5D). Together, these observations support the notion that normal pneumococcal growth requires the synthesis of GTP and/or UTP by *Sp*PYK, pointing to a crucial role of *Sp*PYK in maintaining nucleotide homeostasis.

## DISCUSSION

As one of the model enzymes for studying allosteric regulation, PYK has long been subjected to various structural studies aimed at identifying allosteric sites and characterizing the conformational changes associated with effector binding that alter its kinetic properties. Nevertheless, a detailed understanding of its nucleotide recognition mechanism has been hampered by the dearth of nucleotide-bound PYK structures, which may reflect experimental challenges associated with obtaining high-quality PYK crystals in the presence of nucleotides. It has been speculated that the absence of interactions between the nucleotide base and the active site residues may confer the broad nucleotide specificity observed in PYKs (14). However, the crystal structures reported here show that this does not apply to *Sp*PYK, which can distinguish incoming nucleotides via specific interactions with the nucleotide base. Remarkably, these interactions are not limited to direct interactions between the nucleotide and the protein but also include those mediated by water molecules in the active site pocket, which allow plasticity in the recognition of purine and pyrimidine bases. The E(Q/H)xxR motif on the A172 helix is a key element in nucleotide base recognition, and we anticipate that this newly identified motif could be used to identify PYKs with GDP preference.

An outstanding question relates to the significance of PYK nucleotide selectivity in cell proliferation. PYK has mainly been studied in the context of maintaining glycolytic flux via the conversion of PEP to pyruvate, while its NTP synthesis activity has received less attention (26, 27). However, as one of the two glycolytic enzymes capable of NTP synthesis, PYK plays a prominent role in nucleotide metabolism of *S. pneumoniae*, which is a strictly fermentative bacterium that lacks the tricarboxylic acid cycle and the respiratory electron transport chain (28, 29). To ensure that optimal amounts of NTPs are present throughout the cell cycle, PYK likely evolved to coordinate NTP synthesis with other NTP synthesis pathways. This may explain why CTP synthesis is the least favored reaction by *Sp*PYK because *S. pneumoniae* possesses a dedicated CTP synthase encoded by the *pyrG* gene (30). On the other hand, the absence of such an enzyme for GTP synthesis, together with our kinetics data, makes *Sp*PYK the prime candidate responsible for generating GTP in pneumococcal cells. This is supported by the severe growth defect displayed by AAA mutant cells, which indicates that other putative NTP synthases such as nucleoside diphosphate kinase and adenylate kinase are unable to fully rescue the disruption of nucleotide homeostasis caused by PYK malfunction (31–33).

If GTP is the primary nucleotide product of *Sp*PYK in cells, it is tempting to speculate that GTP plays a more consequential role as an energy source in *S. pneumoniae* compared to other species that can generate large amounts of ATP through oxidative phosphorylation. Although GTP is known to serve as the energy donor in several cellular processes including translation and cell division, ATP is considered to be the universal energy currency (34, 35). This is thought to be the result of an evolutionary pressure arising from the relative intracellular abundance of ATP that shaped enzymes to preferentially use ATP (36). However, the intracellular concentrations of ATP and GTP are comparable in *S. pneumoniae* across different growth phases, which may have relieved this evolutionary pressure to fully commit to using ATP as their primary energy source (37). An intriguing example is the pneumococcal multidrug efflux pump PatAB, which is an ABC transporter that hydrolyzes GTP more efficiently than ATP, unlike other members of this transporter family (37). It is possible that other pneumococcal proteins share this feature, implying the effect of PYK nucleotide selectivity in determining the mode of cellular energy consumption in some species.

In addition to elucidating the physiological role of *Sp*PYK, our study paves a path toward developing specific inhibitors that target a subset of PYKs. This is especially important for antibiotic development that requires selective inhibition of prokaryotic enzymes with high structural homology to their eukaryotic homologs in order to avoid host toxicity (38). *Sp*PYK is an attractive target for drug development because of its essentiality in pneumococcal growth, but no inhibitor has been identified to date. One strategy to selectively inhibit prokaryotic PYKs is to discover compounds that bind to regions distal to the active site like those identified for *Staphylococcus aureus* PYK (39). However, this work suggests that small molecules that interfere with nucleotide base recognition would also be suitable lead compounds for drug development. These molecules could be optimized from previously reported active site binders or antiviral nucleoside analogs containing a guanine backbone (40–42). The wealth of biochemical and structural information on *Sp*PYK now makes it possible to rationally design inhibitors through *in silico* and medicinal chemistry approaches in combination with *in vitro* and cellular validation of target inhibition.

## EXPERIMENTAL PROCEDURES

### Materials and bacterial culture conditions

Culture media were purchased from Becton Dickinson. *E. coli* strains were grown with shaking at 37 °C in lysogeny broth or on agarized lysogeny broth plates. *S. pneumoniae* strains were cultured statically in Todd Hewitt broth containing 0.5% yeast extract (THY) at 37 °C in an atmosphere containing 5% CO_2_. When growth on solid media was required for *S. pneumoniae* strains, pre-poured Trypticase Soy Agar with 5% Sheep Blood (TSAII 5%SB) plates with a 5 mL overlay of 1% nutrient broth agar or TSA plates containing 5% defibrinated sheep blood (Japan BioSerum) with appropriate additives were used. The following concentration of antibiotics were used: carbenicillin, 50 µg/mL; erythromycin, 0.2 µg/mL; kanamycin, 50 µg/mL (*E. coli*) or 250 µg/mL (*S. pneumoniae*); spectinomycin, 200 µg/mL. The bacterial strains, plasmids and oligonucleotide primers used in this study as well as the protocol for plasmid can be found in the Supplementary Information.

### *S. pneumoniae* strain construction

A previously described protocol was used to transform *S. pneumoniae* (43). In brief, upon reaching mid-log phase, *S. pneumoniae* cells were diluted to OD_600_ ∼0.03 in THY containing 1 mM CaCl_2_ and 0.2% bovine serum albumin, and competence was induced by adding 500 ng/mL competence-stimulating peptide (CSP-1; GenScript). After 15 min incubation, 200 ng DNA product was mixed with 1 mL culture, and the resulting culture was further incubated for 1 h before combining it with 5 mL molten 1% nutrient broth agar supplemented with appropriate additives. This mixture was poured onto a TSAII 5%SB plate, and the plate was incubated overnight to recover transformants. Detailed protocols for strain construction can be found in the Supporting Information.

### *S. pneumoniae* fosfomycin sensitivity assay, growth assessment and morphological analysis

Protocols for the fosfomycin sensitivity assay and growth assessment of *S. pneumoniae* cells were adapted from a previous report (18). For determining fosfomycin sensitivity on solid media, *S. pneumoniae* cells at OD_600_ ∼0.2 were collected by centrifugation and normalized to OD_600_ = 1. The normalized samples were diluted 10-fold four times, and 3 µL dilution sample was spotted onto TSA 5% SB plates with or without 64 µg/mL fosfomycin. Plates were imaged after overnight incubation. Growth of *S. pneumoniae* cells was assessed by diluting cultures grown to OD_600_ ∼0.2 to OD_600_ = 0.01 and measuring OD_600_ at the indicated time points. For phase contrast microscopy, live cells were imaged using a Nikon Ts2R inverted microscope equipped with a CFI Plan Fluor 100x 1.3 Oil Ph3 DLL objective and a Nikon Ds-Fi3 camera. A total of 300 non-chained cells and chained cells with widths at constriction sites below 50% of cell widths from two independent experiments were measured for each strain using MicrobeJ (v 5.13o) to determine the cell length and width (44).

### Expression and purification of *Sp*PYK

*Sp*PYK was expressed and purified according to a previously published method with some modifications (18). E*. coli* ECOS Sonic cells harboring the expression plasmid were grown in 1 L lysogeny broth supplemented with kanamycin at 37 °C with shaking until OD_600_ ∼0.4. The culture was cooled to 18 °C and protein expression was induced by adding 500 µM IPTG. Cells were harvested 18 h post-induction by centrifugation (4,200 x g, 15 min, 4 °C). All purification steps were performed at 4 °C. Cells were resuspended in 25 mL HBS (20 mM HEPES pH 7.5, 150 mM NaCl) supplemented with 0.25 mg/mL DNaseI and 0.5 mg/mL lysozyme, and the sample was lysed with French press. Cell debris was removed by centrifugation (10,000 x g, 5 min, 4 °C) and the soluble fraction was collected by ultracentrifugation (100,000 x g, 30 min, 4 °C). The resulting supernatant was supplemented with 0.3 mL pre-equilibrated His60 Ni superflow resin (Takara Bio) and 20 mM imidazole and the resulting mixture was stirred for 30 min at 4 °C. The sample was then loaded onto a gravity column and washed with 30 mL HBS containing 25 mM imidazole followed by 30 mL HBS containing 50 mM imidazole. The protein was then eluted in 3 mL buffer A containing 300 mM imidazole before applying the eluate onto Econo-Pac 10DG column (Bio-Rad) equilibrated with HBS to remove imidazole. Samples were concentrated by centrifugal filtration with an Amicon Ultra-4 centrifugal concentrator (100 kDa cutoff) and immediately used for crystallization trials or flash-frozen with liquid nitrogen and stored at -80 °C for biochemical studies.

### Crystallography and data collection

Vapor diffusion crystallization trials were performed at 25 °C by mixing 15-20 mg/mL protein solution with an equal volume of reservoir solution. The protein solution was prepared by mixing the purified protein sample with KCl, MgCl_2_, oxalate, FBP and nucleotide stocks prepared in HBS (final buffer composition: 20 mM HEPES pH 7.5, 150 mM NaCl, 80 mM KCl, 20 mM MgCl_2_, 20 mM FBP, 20 mM oxalate and 10-20 mM nucleotide). Crystal growth was observed within 4 days, and diffraction-quality crystals were harvested over the course of 1-4 weeks. For cryoprotection, crystals were soaked in reservoir solution containing 20% glycerol (ADP and GDP) or ethylene glycol (ATP and UDP) as cryoprotectant for several minutes. The following reservoir solutions were used to obtain crystals for each structure: ATP — 200 mM potassium acetate and 20% polyethylene glycol (PEG) 3350; ADP and GDP— 100 mM Tricine pH 8.0, 350 mM NaCl, 10% glycerol and 28% PEG 1000; UDP-1 — 20 mM Tris pH 7.0, 100 mM potassium chloride and 20% PEG 4000; UDP-2 — 100 mM HEPES pH 7.2, 150 mM sodium formate and 18% PEG 3350.

Each X-ray diffraction dataset was collected at beamline BL44XU (SPring-8), with a EIGER X 16M detector at 100K. The X-ray diffraction data were processed and scaled using XDS (45). The initial model of PYK was determined by the molecular replacement method using MOLREP in the CCP4 program suite with PEP/FBP-bound *Sp*PYK (PDB: 8IAX) as the structural template (46). The initial model was refined and rebuilt using REFMAC5 and Coot (47, 48). Data collection and refinement statistics are summarized in Table S1. Figures were prepared using PyMOL (v.2.5.8).

### *In vitro* measurement of *Sp*PYK activity

A published protocol with some modifications was used to assess *Sp*PYK activity *in vitro* (18). Measurements were performed at 25 °C in 100 µL reaction mixture containing 1x reaction buffer (20 mM HEPES pH 7.5, 100 mM KCl, 10 mM MgCl_2_, 2 mM FBP, 0.6 mM NADH and 5 U lactate dehydrogenase) and *Sp*PYK. To measure enzyme kinetics, NDP was added to the reaction mixture and the resulting solution was incubated for 10 min before adding 3 mM PEP to initiate reaction. *Sp*PYK activity was assessed by measuring the decrease in absorbance at 340 nm using an Infinite M200 Pro plate reader (Tecan). Kinetic analysis was performed with GraphPad Prism (v.9.4.1) using the Michaelis-Menten model.

## Data availability and accession numbers

X-ray crystallography data have been deposited in Protein Data Bank under the accession numbers 8XW6 (ATP), 8XW7 (ADP), 8XW8 (GDP), 8XW9 (UDP-1), and 8ZLY (UDP-2). All other study data are included within the manuscript.

## Acknowledgements

We thank Yumiko Takeshita for technical assistance with the experiments. X-ray diffraction data were collected using the Osaka University synchrotron beamline BL44XU at SPring-8 (Harima, Japan) under the Collaborative Research Program of Institute for Protein Research (Proposal No. 2021B6624, 2022A6720, and 2022B6720). Funding for this work was provided by the Takeda Science Foundation, Institute for Fermentation (Osaka), Nippon Foundation-Osaka University Project for Infectious Disease Prevention, AMED grant JP23nk0101653, JSPS-NUS Joint Research Program grant JPJSBP120229003 and JSPS KAKENHI grants 21K20759, 23H02631 and 23K14518.

