## Supplementary Information for "Structural basis of nucleotide selectivity in pyruvate kinase"

**Supplementary Methods**

**Plasmid construction**

His-*Sp*PYK variants: Plasmids expressing mutant *Sp*PYK were constructed by introducing the desired mutation in pATOS125 (His-*Sp*PYK) with InFusion Cloning using the following primer pairs: E64A, oAT301/oAT302; Q65A, oAT303/oAT304; R68A, oAT305/oAT306; E64AR68A, oAT307/oAT308; E64AQ65AR68A(AAA), oAT309/oAT310.

***S. pneumoniae* strain construction**

AT1546 (D39 ∆*spd_1735*::*kan*) and AT1553 (D39 ∆*spd_1735*::*kan*, *pyk* (E64A, Q65A, R68A)): A plasmid (pATOS133) encoding a constitutively expressing *pyk* in the absence of LacI was transformed into D39 WT. This strain was transformed with a PCR product containing a *sacBerm* cassette flanked by ~1kb upstream and downstream regions of *pyk* amplified from AT1275 (Taguchi ref) with primers oAT311/oAT312, and the resulting transformant was selected with erythromycin to generate AT1542 (D39 ∆*spd_1735*::(P_lac_-*pyk*); ∆*pyk*::(*erm*, *sacB*)). Primers oAT313/oAT314 were used to amplify *pyk* and *pyk* (E64A, Q65A, R68A) from pATOS125 and pATOS244, respectively, and the amplified products were ligated to upstream and downstream *pyk* regions prepared by PCR using primer pairs oAT311/oAT315 and oAT312/oAT316. AT1542 was then transformed with the resulting PCR products to replace the *sacBerm* cassette, and transformants were selected with 10% sucrose. Integration of PCR cassettes into the *pyk* locus was confirmed via diagnostic PCR using primers oAT317/oAT318 and DNA sequencing. To delete P_lac_-*pyk* integrated in the ∆*spd_1735* locus, a *kan* cassette (oAT319/oAT320) flanked by ~1kb upstream and downstream regions of ∆*spd_1735* prepared by PCR with primer pairs oAT321/oAT322 and oAT323/oAT324 were transformed to the respective strains and transformants were selected with kanamycin to generate AT1546 and AT1553. Whole genome sequencing was performed for AT1553 to confirm that this strain contained the desired replacements without introducing background mutations.

AT1596 (R6 *lacI*, P_lac_-*pyk*, *pyk* (E64A, Q65A, R68A)): The PCR cassette containing the *pyk* (E64A, Q65A, R68A) gene flanked by the *pyk* upstream and downstream regions was transformed into AT1275 (*lacI*, P_lac_-*pyk*, ∆*pyk::erm*, *sacB*) and transformants were selected in the presence of 1 mM IPTG and 10% sucrose. Replacement of the *pyk* locus was confirmed via diagnostic PCR using primers oAT317/oAT318 and DNA sequencing.

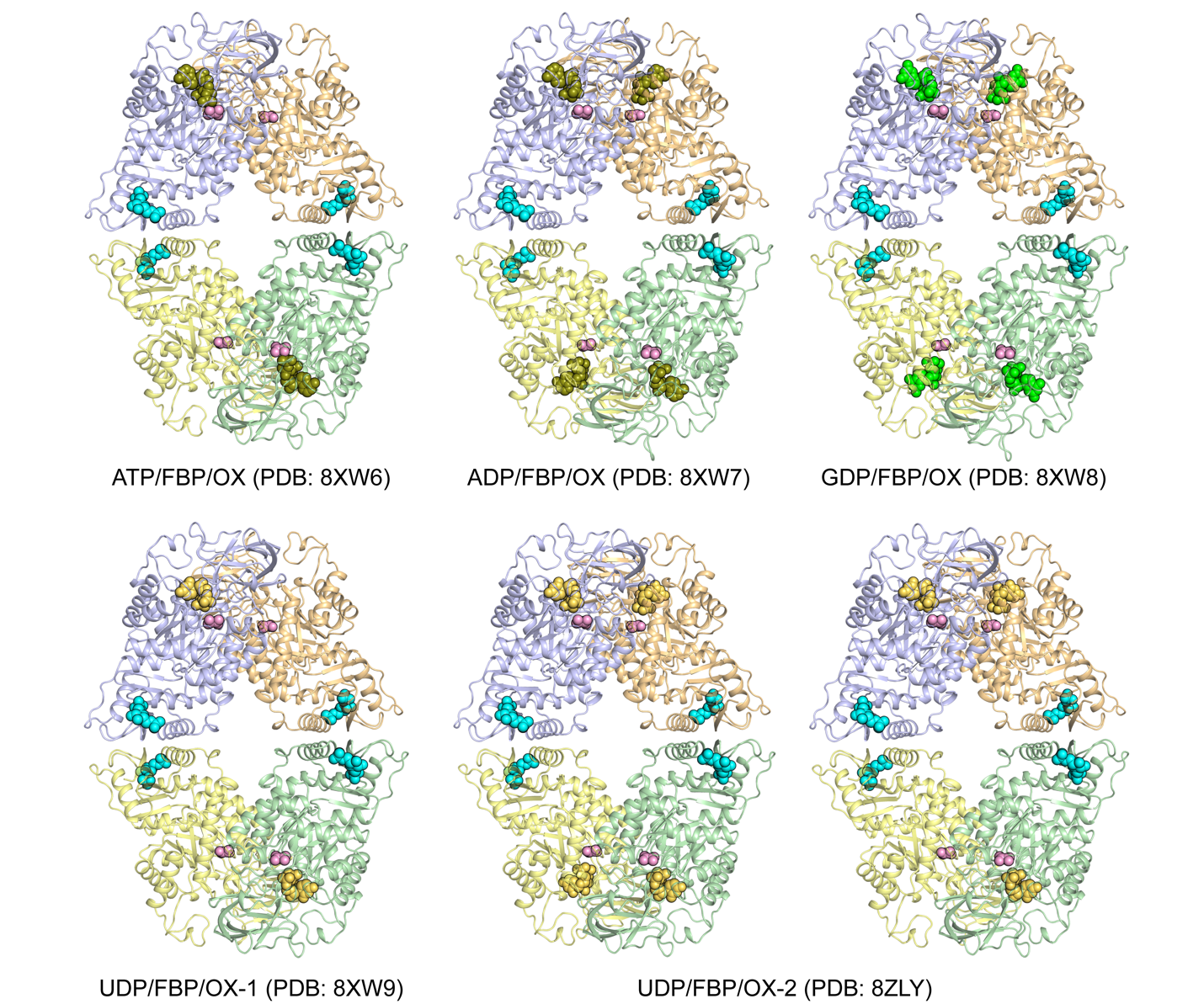

**Figure S1.** **Tetrameric structures of nucleotide-bound *Sp*PYK solved in this study.** ATP/ADP, GDP and UDP in the active site are shown in olive, green, and yellow, respectively. The ATP/FBP/OX and UDP/FBP/OX-1 structures contained two nucleotide molecules per tetramer. Tetramers with three or four UDP molecules were present in the UDP/FBP/OX-2 structure. Oxalate (OX; pink) and FBP (teal) are present in every protomer. Cations are not shown for clarity.

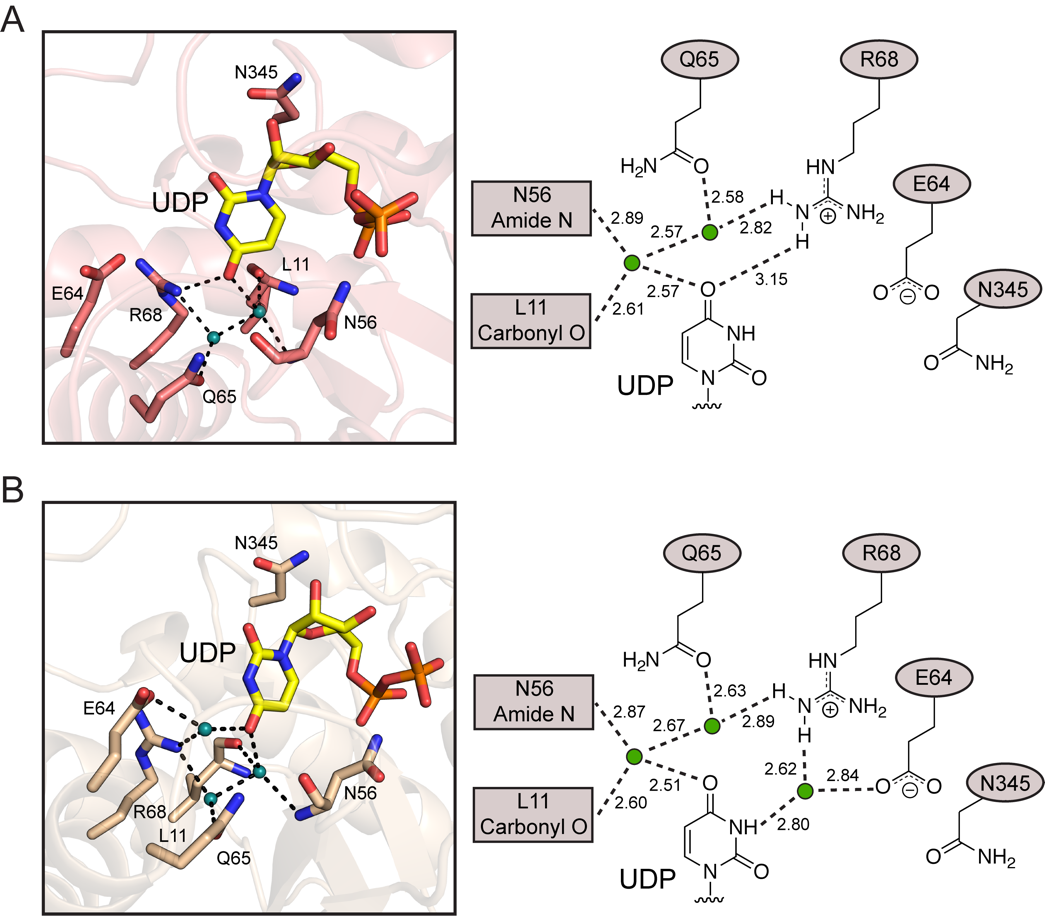

**Figure S2. UDP-bound protomers display differences in how active site residues engage with uridine.** Putative interactions observed in the active site of protomers located in chain C (A) and chain D (B) of the UDP/FBP/OX-2 structure (PDB: 8ZLY) are shown along with the observed distances in angstroms. Teal spheres indicate water molecules. In both protomers, the water molecule that bridges uridine and Gln345 seen in the UDP/FBP/OX-1 structure is missing. In chain C, neither Glu64 nor Asn345 participates in UDP recognition. In chain D, Arg68 interacts with the water molecule that bridges Glu64 and uridine instead of forming a direct interaction with the O4-uridine.

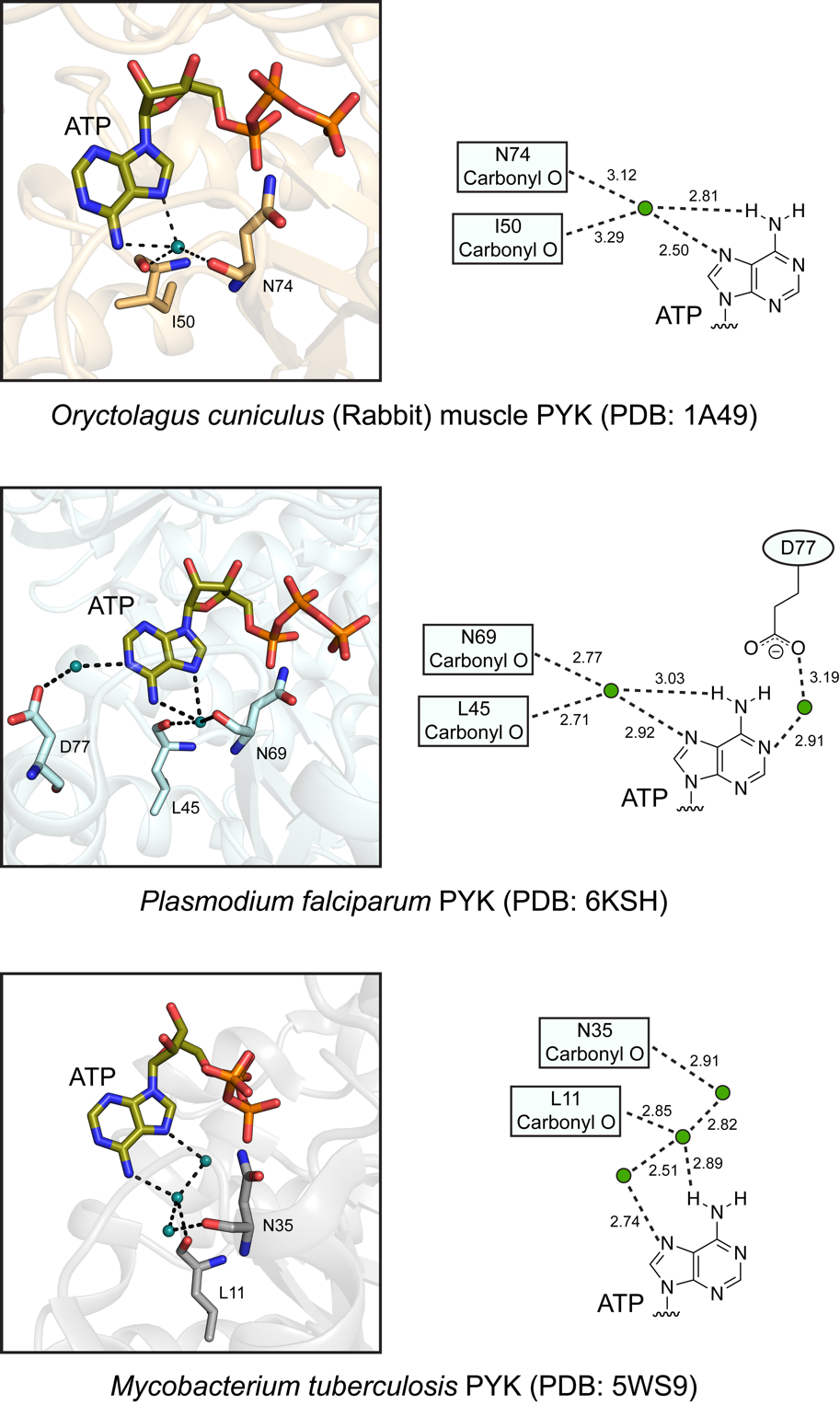

**Figure S3. ATP is coordinated via water molecules in published ATP-bound PYK structures.** Putative interactions between the adenine base and active site residues for rabbit, *P. falciparum*, and *M. tuberculosis* PYKs are shown along with the observed distances in angstroms (1–3). Water molecules are shown as teal spheres.

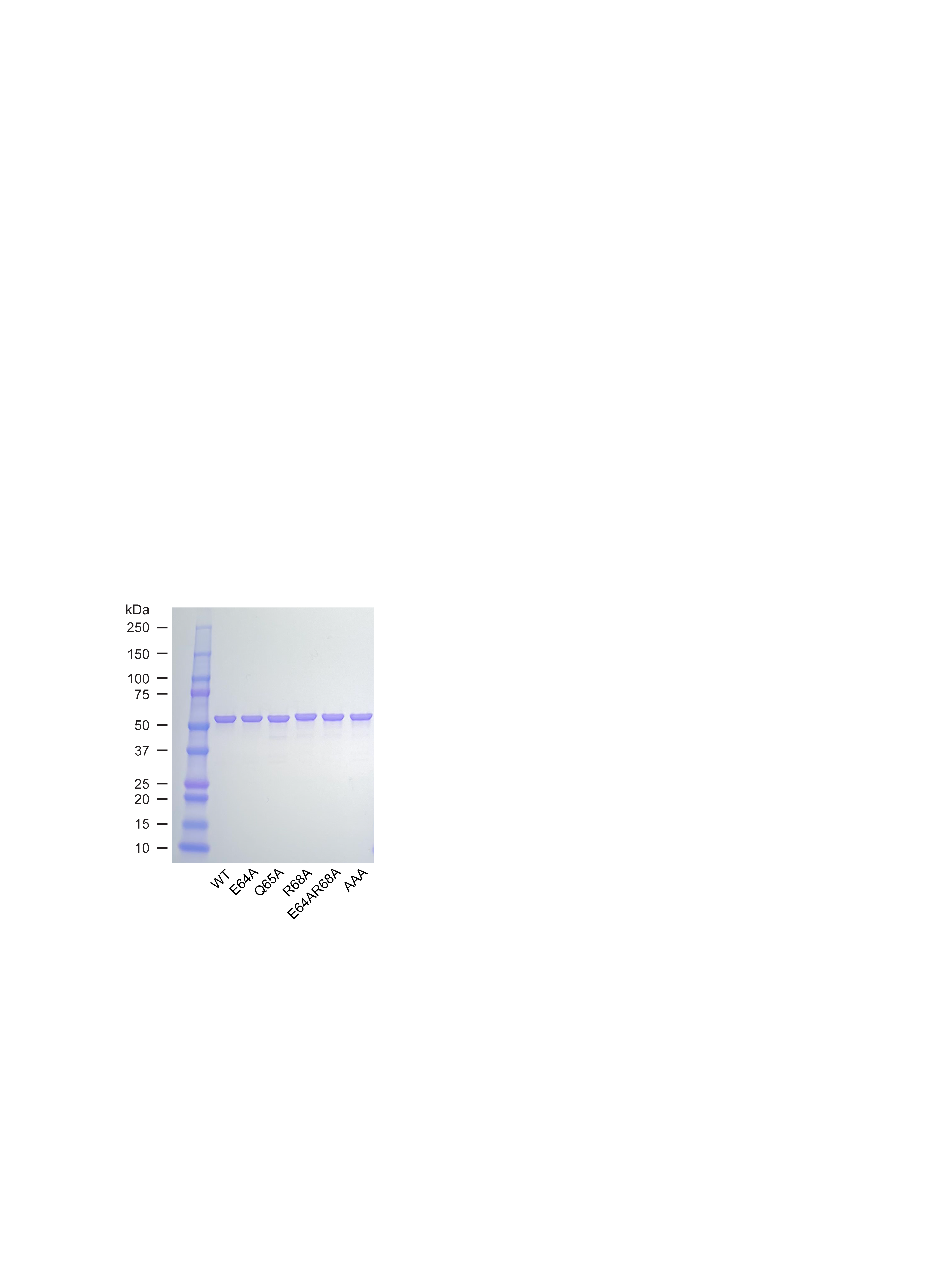

**Figure S4. Coomassie stained gel of purified *Sp*PYK used in this study.** ~2 µg protein was loaded per lane.

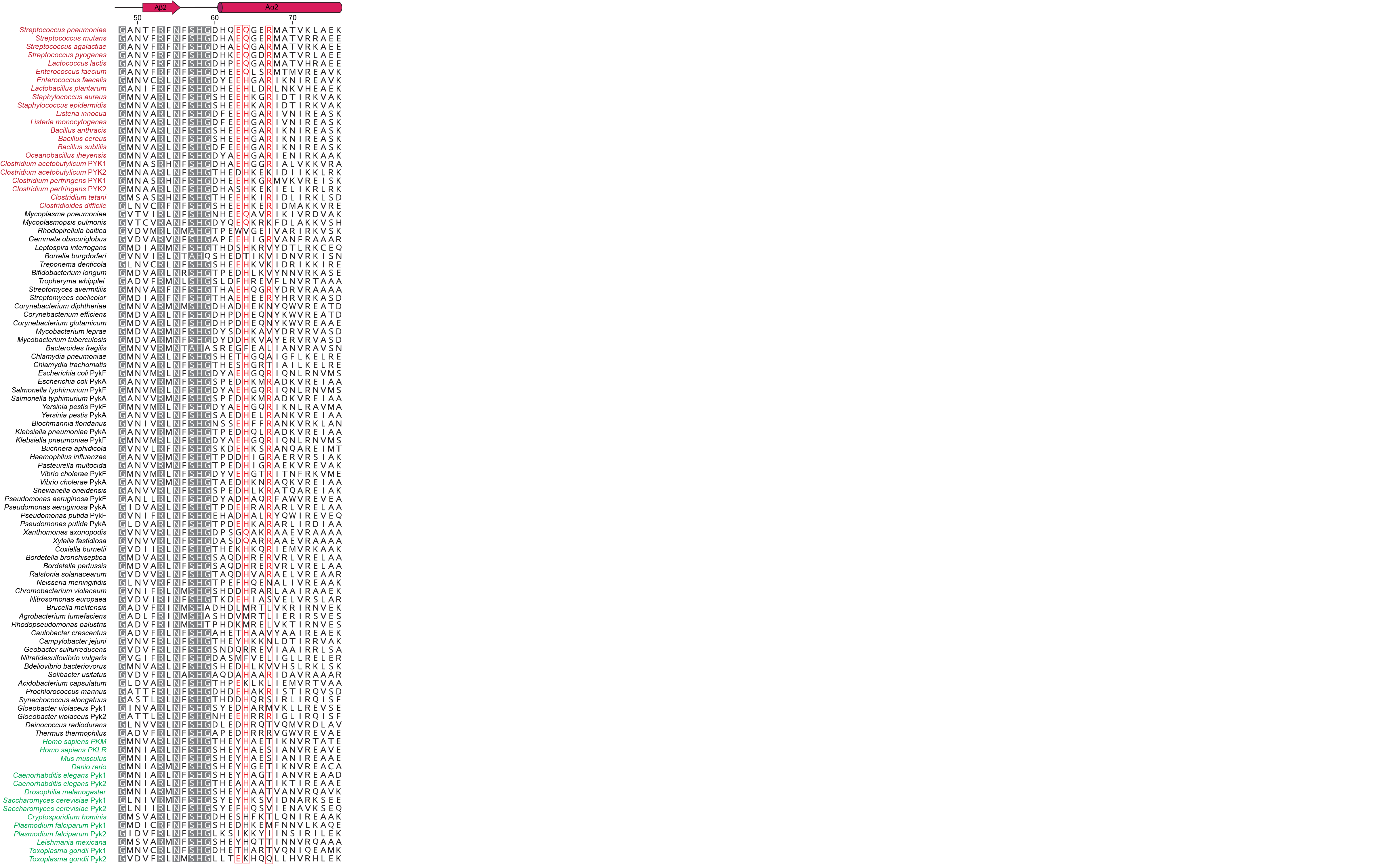

**Figure S5.** **Sequence alignment of the PYK nucleotide recognition domain.** A sequence alignment of 100 PYK sequences from representative prokaryotic and eukaryotic species is shown (Clustal Omega, Geneious Prime v2024.0.5). Secondary structures and residue numbers corresponding to *S. pneumoniae* PYK are shown on the top. Grey residues are conserved in >80% of the PYK sequences that were analyzed in a previous study (4). Firmicutes and eukaryotes are colored in brown and green, respectively. The E(Q/H)xxR motif identified in this study is colored in red. This motif was widely conserved in Firmicutes (colored in brown) and was also found in some Proteobacteria that possess multiple PYK isozymes (e.g. *E. coli*, *K. pneumoniae* and *P. aeruginosa*).

**Table S1. Data collection and refinement statistics for *Sp*PYK.**

|  | **ATP/FBP/OX** | | **ADP/FBP/OX** | **GDP/FBP/OX** | **UDP/FBP/OX-1** | **UDP/FBP/OX-2** |
| --- | --- | --- | --- | --- | --- | --- |
| PDB ID | | 8XW6 | 8XW7 | 8XW8 | 8XW9 | 8ZLY |
| **Data collection** | |  |  |  |  |  |
| Light source | | BL44XU  Spring-8 | BL44XU  Spring-8 | BL44XU  Spring-8 | BL44XU  Spring-8 | BL44XU  Spring-8 |
| Space group | | *P*4_3_22 | *C*222_1_ | *C*2 | *P*4_3_22 | *P*2_1_2_1_2 |
| Wavelength (Å) | | 0.900 | 0.900 | 0.900 | 0.900 | 0.900 |
| Cell dimensions | |  |  |  |  |  |
| *a, b, c* (Å) | | 81.58 81.58 402 | 123.18 255.54 88.133 | 123.33 256.08 88.575 | 81.464 81.464 398.91 | 115.81 403.2 113.59 |
| α, β,γ (°) | | 90.00 90.00 90.00 | 90.00 90.00 90.00 | 90.00 90.041 90.00 | 90.00 90.00 90.00 | 90.00 90.00 90.00 |
| Resolution (Å) | | 2.00 (2.12-2.00) | 2.10 (2.22-2.10) | 2.00 (2.12-2.00) | 1.75 (1.85-1.75) | 1.90 (2.00-1.90) |
| *R*_merge_ | | 7.8 (105.7) | 6.7 (93.8) | 11.2 (91.2) | 5.7 (93.6) | 11.7 (90.6) |
| *I*/*σI* | | 13.59 (1.38) | 19.63 (2.33) | 16.79 (3.64) | 16.01 (1.59) | 13.31 (2.12) |
| Completeness (%) | | 99.3 (98.3) | 100 (99.9) | 97.9 (96.6) | 99.5 (99.0) | 99.8 (98.5) |
| CC (1/2) | | 99.9 (71.6) | 100 (89.5) | 99.8 (82.1) | 99.9 (67.2) | 99.8 (87.7) |
| **Refinement** | |  |  |  |  |  |
| Resolution (Å) | | 46.96-2.00 | 47.22-2.10 | 47.30-2.00 | 40.74-1.75 | 50.0-1.89 |
| Number of reflections | |  |  |  |  |  |
| Observed | | 755658 (118632) | 748128 (113913) | 1305074 (203771) | 816234 (130502) | 5654078 (860413) |
| Unique | | 93443 (14917) | 81724 (13044) | 181214 (28831) | 136776 (21665) | 423736 (67145) |
| *R*_work_/*R*_free_ | | 0.1784/0.2111 | 0.1861/0.2268 | 0.1981/0.2336 | 0.1602/0.1906 | 0.1652/0.2047 |
| Number of atoms | |  |  |  |  |  |
| Protein | | 7680 | 7672 | 15360 | 7680 | 30720 |
| Ligand/ion | | 88 | 118 | 264 | 82 | 406 |
| Water | | 509 | 364 | 1806 | 911 | 3849 |
| B-factors | |  |  |  |  |  |
| Protein | | 51.48 | 57.10 | 32.21 | 36.29 | 38.96 |
| Ligand/ion | | 53.08 | 70.09 | 32.44 | 33.64 | 35.72 |
| Water | | 54.66 | 54.24 | 40.49 | 46.66 | 46.92 |
| RMS deviations | |  |  |  |  |  |
| Bond lengths (Å) | | 0.0115 | 0.0100 | 0.0186 | 0.0137 | 0.0113 |
| Bond angles (°) | | 1.7137 | 1.6574 | 1.8324 | 1.7831 | 1.7187 |

^*^ Highest resolution shell is shown in parentheses.

**Table S2. Bacterial strains used in this study**

| **Strain** | **Description**^*^ | **Reference** |
| --- | --- | --- |
| *E. coli* |  |  |
| ECOS Sonic | BL21(DE3) derivative strain for protein production | Nippon Gene |
| *S. pneumoniae* |  |  |
| D39 | Wild-type | S. Kawabata |
| AT1546 | D39 ∆*spd_1735*::*kan*; Kan^R^ | This study |
| AT1553 | D39 *pyk* (E64A, Q65A, R68A), ∆*spd_1735*::*kan*; Kan^R^ | This study |
| R6 | Wild-type | S. Kawabata |
| AT1023 | R6 *∆prsA*::(PF6-*lacI*, *gent*), ∆*spr1750* ::(P_lac_-*pyk*); Gent^R^, Spec^R^ | (4) |
| AT1408 | R6 *∆prsA*::(PF6-*lacI*, *gent*), ∆*spr1750* ::(P_lac_-*pyk*); *pyk* (H411A); Gent^R^, Spec^R^ | (4) |
| AT1596 | R6 *∆prsA*::(PF6-*lacI*, *gent*), ∆*spr1750* ::(P_lac_-*pyk*); *pyk* (E64A, Q65A, R68A); Gent^R^, Spec^R^ | This study |

**Table S3. Plasmids used in this study**

| **Plasmid** | **Description**^*^ | **Reference** |
| --- | --- | --- |
| pATOS125 | His_6_-*pyk* expression vector; Kan^R^ | (4) |
| pATOS235 | His_6_-*pyk* (E64A) expression vector; Kan^R^ | This study |
| pATOS241 | His_6_-*pyk* (Q65A) expression vector; Kan^R^ | This study |
| pATOS242 | His_6_-*pyk* (R68A) expression vector; Kan^R^ | This study |
| pATOS243 | His_6_-*pyk* (E64A, R68A) expression vector; Kan^R^ | This study |
| pATOS244 | His_6_-*pyk* (E64A, Q65A, R68A) expression vector; Kan^R^ | This study |
| pATOS133 | *S. pneumoniae spr1750*::P_lac_-*pyk* integration vector; Spec^R^ | (4) |

^*^Abbreviations: Gent^R^, gentamicin resistance; Kan^R^, kanamycin resistance; Spec^R^, spectinomycin resistance

**Table S4. Oligonucleotide primers used in this study**

| **Primer** | **Sequence (5ʹ-3ʹ)** |
| --- | --- |
| oAT301 | CCACCAAGCACAAGGTGAGCGTATGGCAAC |
| oAT302 | CCTTGTGCTTGGTGGTCGCCGTGTGAG |
| oAT303 | CCAAGAAGCAGGTGAGCGTATGGCAACTGT |
| oAT304 | TCACCTGCTTCTTGGTGGTCGCCGTG |
| oAT305 | AGGTGAGGCTATGGCAACTGTTAAACTTGCGG |
| oAT306 | GCCATAGCCTCACCTTGTTCTTGGTGGTCG |
| oAT307 | AGCACAAGGTGAGGCTATGGCAACTGTTAAACTTGCGGA |
| oAT308 | GCCTCACCTTGTGCTTGGTGGTCGCCGTGTGA |
| oAT309 | AGCAGCAGGTGAGGCTATGGCAACTGTTAAACTTGCGGA |
| oAT310 | GCCTCACCTGCTGCTTGGTGGTCGCCGTGTGA |
| oAT311 | AGAAAAAGGAGCATAAACCAATA |
| oAT312 | TCTCCTAGCGATATCCAATC |
| oAT313 | ATATATAAAGGAGTCACAAAAATCATGAACAAACGTGTAAAAATCGT |
| oAT314 | CAGGCTAGAAAGCTGGATATGATAGGTTTTTATATTTTTCTTAACGTACTGTGCGGATAC |
| oAT315 | GATTTTTGTGACTCCTTTATATAT |
| oAT316 | ATATCCAGCTTTCTAGCCTGTA |
| oAT317 | ACAACCCACACGAAGCCG |
| oAT318 | CCATTATGAAAGTTTTGCCC |
| oAT319 | GAGGGAGGAAAGGCAGGA |
| oAT320 | CGCCGTATCTGTGCTCTC |
| oAT321 | ACGCACACTCAACTGGG |
| oAT322 | TCCTGCCTTTCCTCCCTCAACAGTACTTAAAGCTATCCG |
| oAT323 | GAGAGCACAGATACGGCGCTTCTATATGGTGGTGCTTATGC |
| oAT324 | CACAGGCTCTAATTCTATCAC |
